# Crystal structure of steroid reductase SRD5A reveals conserved steroid reduction mechanism

**DOI:** 10.1101/2020.07.23.217620

**Authors:** Yufei Han, Qian Zhuang, Bo Sun, Wenping Lv, Sheng Wang, Qingjie Xiao, Bin Pang, Youli Zhou, Fuxing Wang, Pengliang Chi, Qisheng Wang, Zhen Li, Lizhe Zhu, Fuping Li, Dong Deng, Ying-Chih Chiang, Zhenfei Li, Ruobing Ren

## Abstract

Steroid hormones are essential in stress response, immune system regulation, and reproduction in mammals. Steroids with 3-oxo-Δ^4^ structure, such as testosterone, androstenedione and progesterone, could be catalyzed by steroid 5α-reductases (SRD5As) to generate their corresponding 3-oxo-5α steroids, which are essential for multiple physiological and pathological processes. Abnormal activities of SRD5As will lead to benign prostatic hyperplasia, alopecia, prostatic cancer or infertility due to the poor quality of sperms. However, the detailed reduction mechanisms of SRD5As remain elusive. Here we report the crystal structure of PbSRD5A, which shares 60.6% and 51.5% sequence similarities with human SRD5A1 and −2 respectively, from *Proteobacteria bacterium* in complex with the cofactor NADPH at 2.0 Å resolution. PbSRD5A exists as a monomer comprised of seven transmembrane segments (TMs). The TM1-4 enclose a hydrophobic cavity for steroids substrates binding, whereas TM5-7 coordinate with cofactor NADPH through extensive hydrogen bonds network. Homology-based structural models of HsSRD5A1 and −2, together with extensive biochemical characterizations, for the first time unveiled the substrate recognition of SRD5As and provide an important framework for further understanding of the mechanism of NADPH mediated steroids 3-oxo-Δ^4^ reduction. Based on these analyses, the design of therapeutic molecules targeting SRD5As with improved specificity and therapeutic efficacy would be possible.

**One Sentence Summary:** Structural and biochemical characterizations decipher the evolutionarily conserved mechanism in steroid 5α-reductases catalyzing NADPH mediated steroids reduction.

---

Steroid hormones, derived from cholesterol through *de novo* steroidogenesis in the adrenal cortex, the gonads and the placenta^1,2,^ are essential in stress response, immune system regulation, and reproduction in mammals^3–5^. Testosterone, progesterone and androstenedione are important steroid hormones. Testosterone could be catalyzed to dihydrotestosterone (DHT) which is more potent to activate androgen receptor (AR)-regulated genes^6–9^. Progesterone, the most important progestogen in human, is involved in multiple physiological processes through different nuclear receptors and GABA receptor^10,11.^ Androstenedione is the precursor of androgen and estrogen^12^. These steroids could be converted to their corresponding 3-oxo-5α steroids by SRD5As for proper physiological function (Fig. 1a)^13,14.^

**Figure 1:**
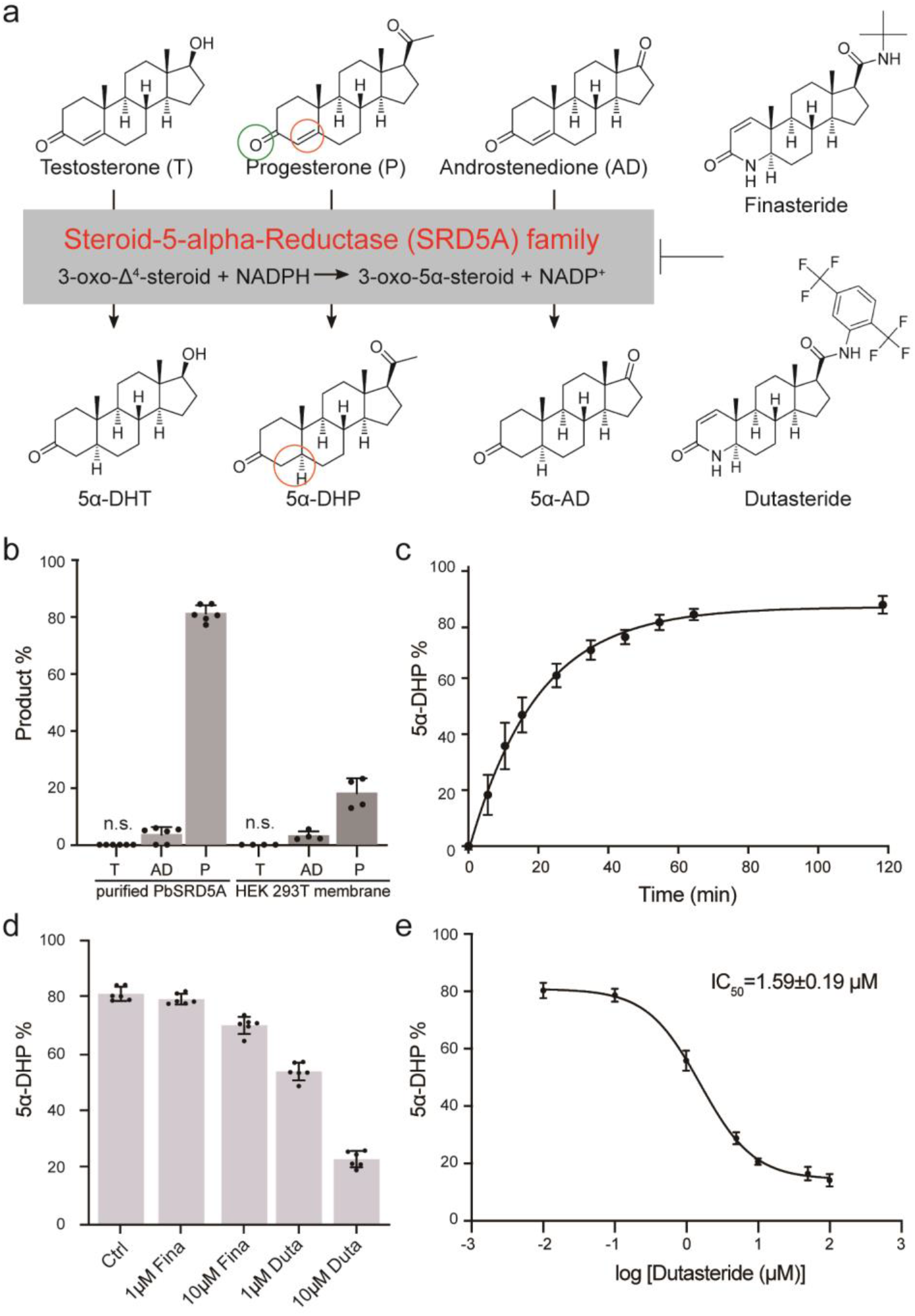
Steroid 5α-reductase activity of PbSRD5A. **a,** Steroid 5α-reductase family proteins can reduce Δ^4^ carbon-carbon double bound to form α configuration at C^5^ of products. These reactions can be inhibited by finasteride and dutasteride specifically. **b,** Testosterone (T), androstenedione (AD) and progesterone (P) metabolism *in vitro* and in HEK 293T cell. Purified PbSRD5A and 293T cells transiently transfected with PbSRD5A were treated with [^3^H]-labeled T, AD and P for 1 h and 12 h, respectively. n.s. means no signal. **c,** Time curve of reduction reaction by PbSRD5A to progesterone. **d,** Inhibition of PbSRD5A by finasteride (Fina) and dutasteride (Duta) *in vitro*, both with the concentration of 1 μM and 10 μM, respectively. **e,** Dutasteride inhibits the activity of PbSRD5A *in vitro*, with an IC50 value of 1.58±0.19 μM (logIC50=0.20±0.05). In **b-e**, the percentages of products were measured using high-performance liquid chromatography (HPLC) with detected radioactivity. Data are mean±s.d. derived from technically independent experiments in duplicate. Each experiment was reproduced at least three times on separate occasions with similar results.

Human steroid 5α-reductase isozymes, belonging to NADPH dependent oxidoreductase family EC1.3.1.22, have three members (SRD5A1-3). SRD5A1 and −2 are mainly involved in the metabolism of steroid hormones^15^, whereas SRD5A3 converts polyprenol into dolichol for the early steps of protein N-linked glycosylation^16,17.^ Both SRD5A1 and −2 are endoplasmic reticulum membrane-embedded and utilize NADPH as electron and proton donor for the Δ^4^ reduction reactions^18–20^. However, SRD5A1 prefers androstenedione as substrate to generate 5α-androstenedione and SRD5A2 prefers testosterone to produce DHT^21,22.^ During prostate cancer progression^23–25^, decreased SRD5A2 activity and increased SRD5A1 activity have been observed and the related clinical significance remains to be explored^26^. Several inhibitors have been developed to inhibit the activity of SRD5A1 and −2 for disease treatment (Fig. 1a)^27,28.^ Finasteride specifically inhibits SRD5A2 and dutasteride inhibits both SRD5A1 and −2^29^, both of which are widely used for benign prostatic hyperplasia (BPH)^30–34^. It is worth mentioning that, recent data around the world indicated men had a greater risk of severe illness and death of COVID-19 than women and children^35^. One of the possible reason is that, androgens such as testosterone appeared to boost the virus’ ability entering into the cell by elevating the expression level of TMPRSS2^36^, which is a well characterized gene regulated by androgens in prostate cancer patients^37^. Androgen-driven immune modulation may also contribute to the COVID-19 associated clinical outcomes^38^. Prostate cancer patients with COVID-19 infection on androgen-deprivation therapy (ADT) were less likely to be hospitalized and to die, compared with those were not on ADT^39^. Cell-based studies also indicated the therapeutic potential of SRD5As’ inhibitor in blocking virus’ entry^40,41.^ Although SRD5A1 and −2 have been identified and extensively explored for decades, the structural information and molecular reaction mechanism are still poorly understood.

## Results

### Functional characterizations of PbSRD5A

To unveil the molecular mechanisms underlying the substrate recognition and catalytic reaction of SRD5A1 and −2, BLAST searches^42^, using HsSRD5A1 and −2 as queries, against the sequenced bacterial genomes were performed to identify a steroid 5α-reductase suitable for structural investigation. Elaborative selection was performed after the identification of hundreds of SRD5A candidates and four bacterial homologs, including PbSRD5A, were cloned, expressed and purified using the insect cell expression system. PbSRD5A shared 60.6% and 51.5% sequence similarities with HsSRD5A1 and −2, respectively. PbSRD5A presented monodisperse behavior on size exclusion chromatography and was selected for the structural and functional characterization due to its higher similarity to HsSRD5As (Extended Data Fig. 1a). To validate the function of PbSRD5A, steroid reduction assays were carried out. PbSRD5A presented potent reduction activity and obvious selectivity to progesterone, traceable activity to androstenedione, but no detectable activity to testosterone, albeit with the similar structures among three steroids (Fig. 1b, c). Interestingly, dutasteride inhibited the reduction reaction of progesterone with a half-maximal inhibitory concentration (IC_50_) of 1.59±0.19 μM (Fig. 1d, e). However, finasteride showed mild inhibition only at high concentration (Fig. 1d).

### Overall architecture of PbSRD5A

After extensive crystallization trails, the crystals of PbSRD5A in space group C2221 yielded in several conditions under monoolein lipid using lipidic cubic phase crystallization method (Extended Data Fig. 1b). The native diffraction data sets were collected at ~2.0 Å resolution. The structural model of PbSRD5A, derived from the modified framework of trRosetta aided by a fused deep residual network that took input multiple sequence alignments, were applied as the initial phase for structure determination (Extended Data Fig. 1c). The structure was determined by molecular replacement using the predicted structural model and refined to 2.0 Å resolution (Fig. 2a, Extended Data Fig. 1d and Table 1).

**Figure 2:**
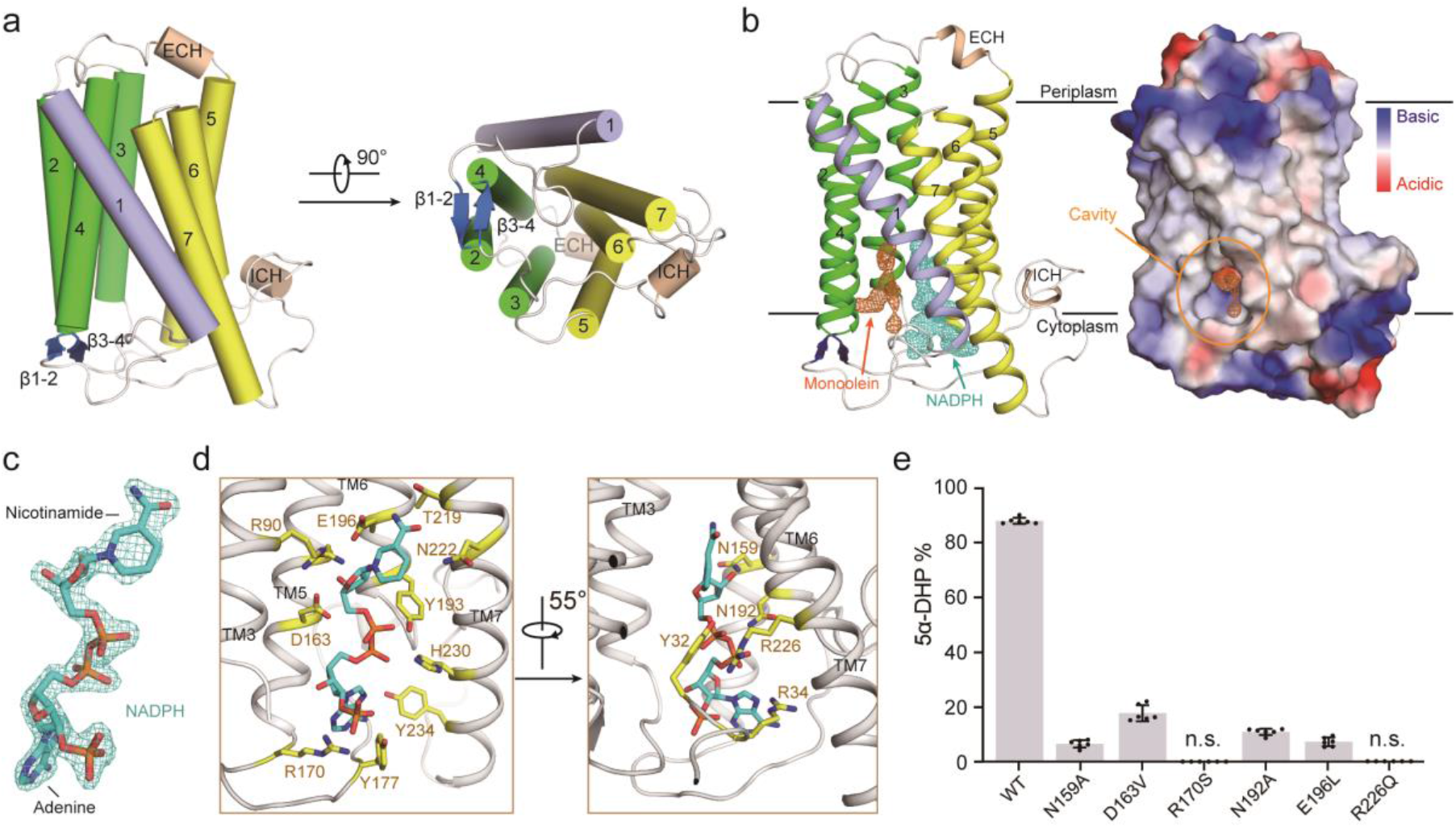
The crystal structure of PbSRD5A and coordination of NADPH. **a,** The overall structure of PbSRD5A. Two perpendicular views are shown. The seven transmembrane segments are divided into TM1 (lightblue), TM2-4 (green), and TM5-7 (yellow). The extracellular and intracellular short alpha helices (ECH and ICH) are colored wheat. The short anti-parallel beta strands are colored blue. The loop regions are colored white. **b,** 2Fo-Fc map for monoolein (orange mesh) and NADPH (cyan mesh) are both contoured at 1.0σ. The two black lines show the approximate location of the lipid bilayer. The cavity is circled in an electrostatic surface representation. **c,** NADPH fits into the density map shown in panel B. Nicotinamide and adenine groups are labeled. **d,** Coordination of NADPH by polar residues of PbSRD5A. The residues that are hydrogen-bonded to NADPH are shown in yellow sticks. **e,** Functional validation of residues coordinating NADPH in PbSRD5A. Enzyme activities were measured by the percentage of 5α-DHP reduced from [^3^H]-labeled progesterone *in vitro* within 1 h detected by HPLC. Data are mean±s.d. derived from technically independent experiments in duplicate. Each experiment was reproduced at least three times on separate occasions with similar results.

PbSRD5A comprises seven transmembrane segments (TMs), with the N-termini located on the periplasmic side of the plasma membrane and C-termini on the cytosolic side. Given the level of sequence conservation, it is likely that all SRD5As exhibit the same fold as PbSRD5A (Extended Data Fig. 1f). We applied Dali search using PbSRD5A structure and found that the TM2-7 of PbSRD5A can be superimposed with TM 5-10 of the integral membrane delta (14)-sterol reductase MaSR1 (Protein Data Bank code: 4QUV) (Extended Data Fig. 2a), but TM2-4 of MaSR1 were missing in PbSRD5A (Extended Data Fig. 2b). Notably, the TM1 conformations of these two proteins were largely different (Extended Data Fig. 2b). The connecting sequence between TM1 and TM2 adapted a short β strand (β1-2), which formed antiparallel β-sheets with the corresponding strand β3-4 linking TM3 and TM4 (Fig. 2a). The linkages between TM4, TM5 and TM6 adapted to short helices and stabilized the protein conformation through hydrogen bonds and hydrophobic interactions with other loops as well as C-termini (Fig. 2a).

There is one PbSRD5A molecule in each asymmetric unit. Examination of the crystal lattice revealed three interfaces, two of which are due to the crystal packing to the neighboring symmetry-related molecules and the third interface is indeterminate (Extended Data Fig. 3a, b). This interface was mediated exclusively through van der Waals interactions between TM1 of two adjacent protomers (Extended Data Fig. 3c). To examine the oligomerization state of PbSRD5A in solution, we applied the PbSRD5A and MvINS (PDB code: 4XU4) to analytical ultracentrifugation (AUC) analysis. The theoretical molecular weight of PbSRD5A and MvINS protomers are 29.0 kDa and 23.6 kDa, respectively. The AUC results indicated that PbSRD5A functioned as a monomer (27.5 kDa) in solution and MvINS exhibited as a trimer (66.6 kDa) (Extended Data Fig. 3d), consistent with the previous cross-linking results^43^.

### Coordination of NADPH

With most amino acids assigned in the electron density map, two omitted electron densities were clearly visible in the central cavity near the cytoplasmic side (Fig. 2b). NADPH perfectly fitted into one of the densities and was majorly coordinated by TM5-7 (Fig. 2b, c). NADPH formed extensive hydrogen bonds with N159, D163 and R170 on TM5, N192, Y193, and E196 on TM6, and T219, N222, R226 and H230 on TM7 (Fig. 2d and Extended Data Fig. 4a, b). The residues on intracellular loop 1 (Y32 and R34) and loop 3 (Y177) also contributed to NADPH binding through direct hydrogen bonds (Fig. 2d and Extended Data Fig. 4a, b). Some residues, such as R34 on intracellular loop 1, V100 on intracellular loop 2, N192 on TM6 and R226 on TM7 partly interacted with NADPH through water mediated hydrogen bonds (Extended Data Fig. 4c). Additionally, the residues W50 on TM2, F94 and M98 on TM3, L166 and L169 on TM 5, and L223 on TM7 interacted with NADPH by hydrophobic effects (Extended Data Fig. 4c). The results of *in vitro* reduction assay demonstrated that mutations in these residues of PbSRD5A impaired the conversion from progesterone to 5α-dihydroprogesterone, indicating the crucial role of these residues for reductase activity (Fig. 2e). The other elongated strip density was modeled with the crystallization lipid monoolein, whose alkenyl “tail” is accommodated well (Fig. 2b and Extended Data Fig. 5a) in the hydrophobic pocket majorly composed by the residues located on TM1-4, such as F17, S21 T24 and L25 on TM1, Y32 on intracellular loop 1, A49 and W50 on TM2, and T109, A110 A113 and F116 on TM4 (Extended Data Fig. 5b). The residue Q53 formed the hydrogen bond with the glycerol “head” of monoolein (Extended Data Fig. 5b). Considering of the hydrophobicity of steroid substrates and the orientation of NADPH, more specifically the position of nicotinamide, we hypothesized that monoolein probably occupied the steroid substrates binding pocket. For clear narration, we named TM1-4 as substrate-binding domain (SBD) and TM5-7 as NADPH binding domain (NDPBD) (Extended Data Fig. 1f).

### Molecular mechanism of 3-oxo-Δ^4^ reduction

Considering the sequence similarities of bacterial SRD5A, HsSRD5A1 and −2, we built the homology models of HsSRD5A1 and −2 using the server for initial model building of PbSRD5A (Fig. 3a, b and Extended Data Fig. 6). The highly conserved residues were mainly concentrated to the NADPH and the putative steroid binding pockets, as well as the interfaces of TMs (Extended Data Fig. 6). It suggested the evolutionary conservation of NADPH binding and substrate recognition other than the overall architectures. The movements of TM1s were observed after we superimposed TM2-7 of PbSRD5A structure with HsSRD5A1 and −2 models (Extended Data Fig. 7a).

**Figure 3:**
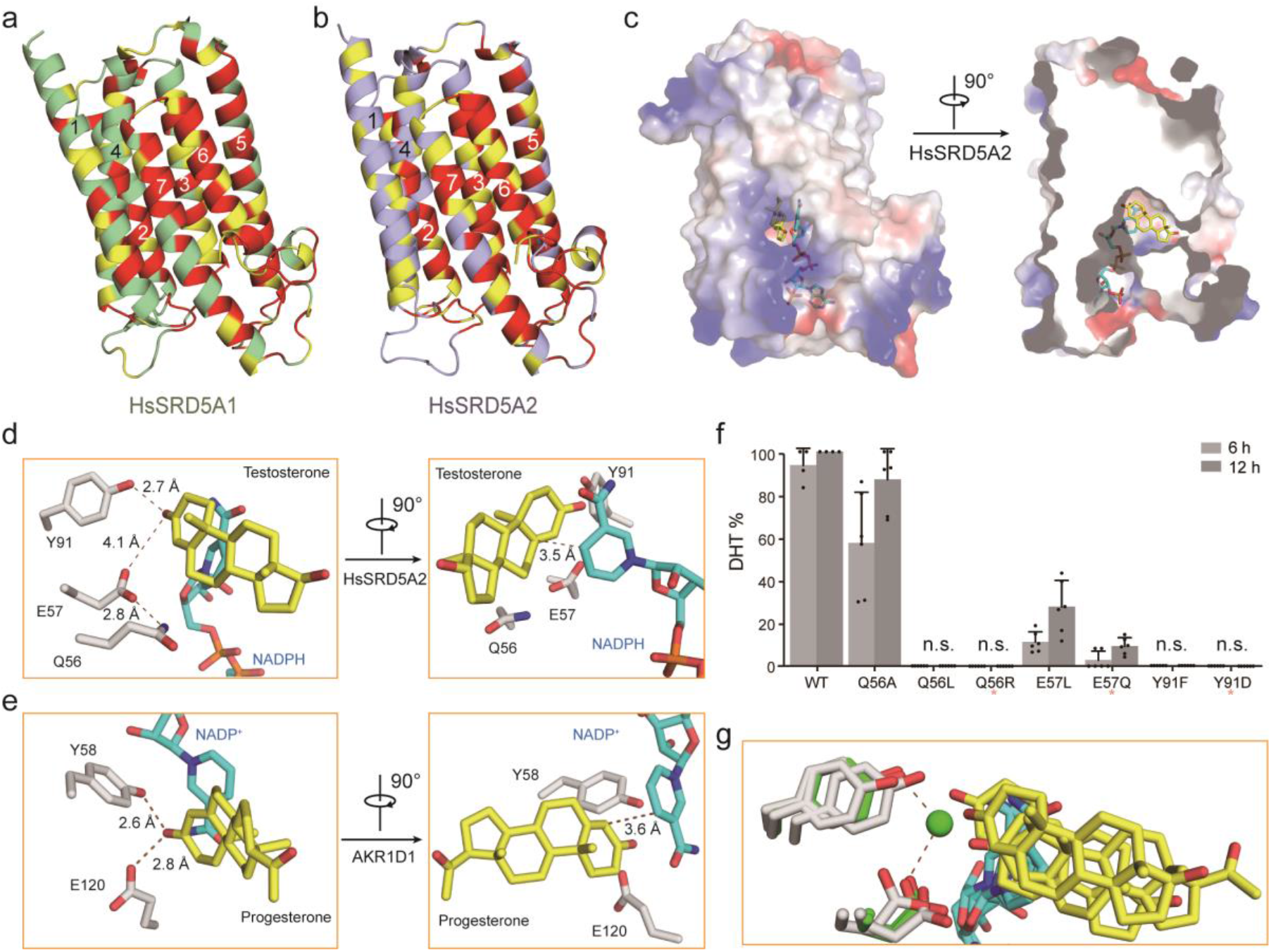
HsSRD5A1 and −2 modeling, substrate docking and functional characterization. **a,** HsSRD5A1 homology model was generated on the basis of PbSRD5A structure and represented in palegreen and lightblue. Invariant and conserved residues are colored in red and yellow, respectively. **b,** HsSRD5A2 homology model was generated on the basis of PbSRD5A structure. **c,** The docking pose of testosterone in HsSRD5A2 structural model. The semi-transparent electrostatic surface of HsSRD5A2 is shown. Testosterone is shown as yellow stick. NADPH is colored cyan. **d,** The coordination of conserved Q-E-Y motif with testosterone in HsSRD5A2 docking model. Two perpendicular views are shown. **e,** The coordination of conserved Q-E-Y motif in AKR1D1-progesterone complex structure. **f,** Biochemical characterization of the Q-E-Y motif of HsSRD5A2. All variants were transient expressed in HEK293T cells for 24 hours. The cells were treated with [3H]-labeled T for 6 or 12 hours. The collected and homogenized cell membranes were treated with [^3^H]-labeled T. HsSRD5A2 activities were measured by the percentage of DHT detected by HPLC. Data are mean±s.d. derived from technically independent experiments in duplicate. Each experiment was reproduced at least three times on separate occasions with similar results. Red stars (*) indicated the variants in the–patients. **g,** The water molecule, coordinated by the Q-E-Y motif in PbSRD5A structure, was shown in green sphere. The water molecule occupied the space for the C-3 carbonyl oxygen in the docking results shown in **Fig. 3d** and **Extended Data Fig. 7d, f.**

To fully explore the substrate recognition and reduction reaction mechanism, progesterone was docked into PbSRD5A, as well as testosterone into HsSRD5A1 and −2, respectively. For instance, in the SRD5A2 docking model, the native substrate testosterone can be accommodated in the semi-closed pocket composed by SBD (TM1-4), which is similar to that in PbSRD5A occupied by monoolein (Fig. 2b and 3c). This binding pose was also observed in the catalytic pocket of soluble steroid 5β-reductase3 (AKR1D1, Protein Data Bank code: 3COT), bound with progesterone (Extended Data Fig. 7b)^44^. Notably, the steroid binding pockets of both enzymes contain a signature motif forming triangular hydrogen bonds that coordinate the ketone group at C-3 in the steroids (Fig. 3d, e). This coordination brings the substrate into close proximity to the nicotinamide of NADPH, leading to the hydride (H^-^) transfer and Δ^4^ double bond reduction (Fig. 3d, e). For HsSRD5A2, this signature motif includes Y91 and E57 (Fig. 3d) and for AKR1D1, Y58 and E120 (Fig. 3e)^44^. Additionally, Q56 of HsSRD5A2 bonded with E57 to stabilize the local conformation in HsSRD5A2 (Fig. 3d). In the reductase activity assay using cells containing HsSRD5A2 mutants, Q56A/L, E57L and Y91F lost reductase activity unambiguously (Fig. 3f), indicating the importance of these residues in substrate recognition and reduction. The Q56R, E57Q and Y91D variants, whose activities were also largely aborted in the reductase assay (Fig. 3f), were found in SRD5A2 deficient disease patients, such as Pseudovaginal perineoscrotal hypospadias (PPSH) ^45–48^. The “QEY” signature motif is highly conserved in all 5α-reductases we studied, including PbSRD5A and HsSRD5A1, demonstrating the evolutionarily conserved steroid reduction mechanism (Extended Data Fig. 7c-f). In the substrate-free PbSRD5A structure, Y87 donates a hydrogen bond to a water molecule, which in turn hydrogen bonded with E54. Interestingly, this hydrogen-bonded water molecule occupies a position close to that of carbonyl oxygen of substrates observed in all the docking models, which is consistent with the observation in AKR1D1 structures (Fig. 3g)^44^.

### Disease related mutagenesis of HsSRD5A2

Due to the essential function of SRD5A2 in steroidogenesis, large numbers of disease-related loss-of-function mutations have been identified and confirmed *in vivo*^49,50.^ Guided by a homologous structural model of HsSRD5A2, we classified 32 mutations into 5 categories (Extended Data Table 2) and mapped on the HsSRD5A2 structural model (Fig. 4a). As discussed previously, the residues Q56, E57 and Y91 colored in magenta are located in the catalytic pocket and essential for the substrate binding and catalytic activity (Fig. 4a left). The residues for NADPH binding, including N160, D164, R171, N193, E197, R227, H231 and Y235, are colored in yellow (Fig. 4a left). The corresponding residues in PbSRD5A examined by *in vitro* reduction assay are all identical to HsSRD5A2 (Fig. 2e and Extended Data Table 2). The residues which form extensive hydrogen bond network to stabilize the protein conformation, but not directly interact with substrates and NADPH, are also highlighted (Fig. 4a middle). Notably, the variants P181L, S245Y and R246Q/W in the C-termini are also disease-related, indicating the C-terminal local conformation is crucial for reductase activity. Considering these residues are close to the NADPH binding site, these variants may destabilize the NADPH binding (Fig. 4a middle). Besides, numbers of Gly to X (colored in wheat) and X to Gly (colored in cyan) variants are found in PPSH patients (Fig. 4a right). These mutations are mainly located on the TM interfaces and may reduce the HsSRD5A2 activity by disrupting the tertiary structure. The disease-related mutants are also extensively studied by steroid reduction assay (Fig. 4b). Except for H231A, the activities of all other variants decreased significantly. The structural analysis combining with biochemical studies clearly elucidated how disease-related mutations abort HsSRD5A reductase activity at the molecular level.

**Figure 4:**
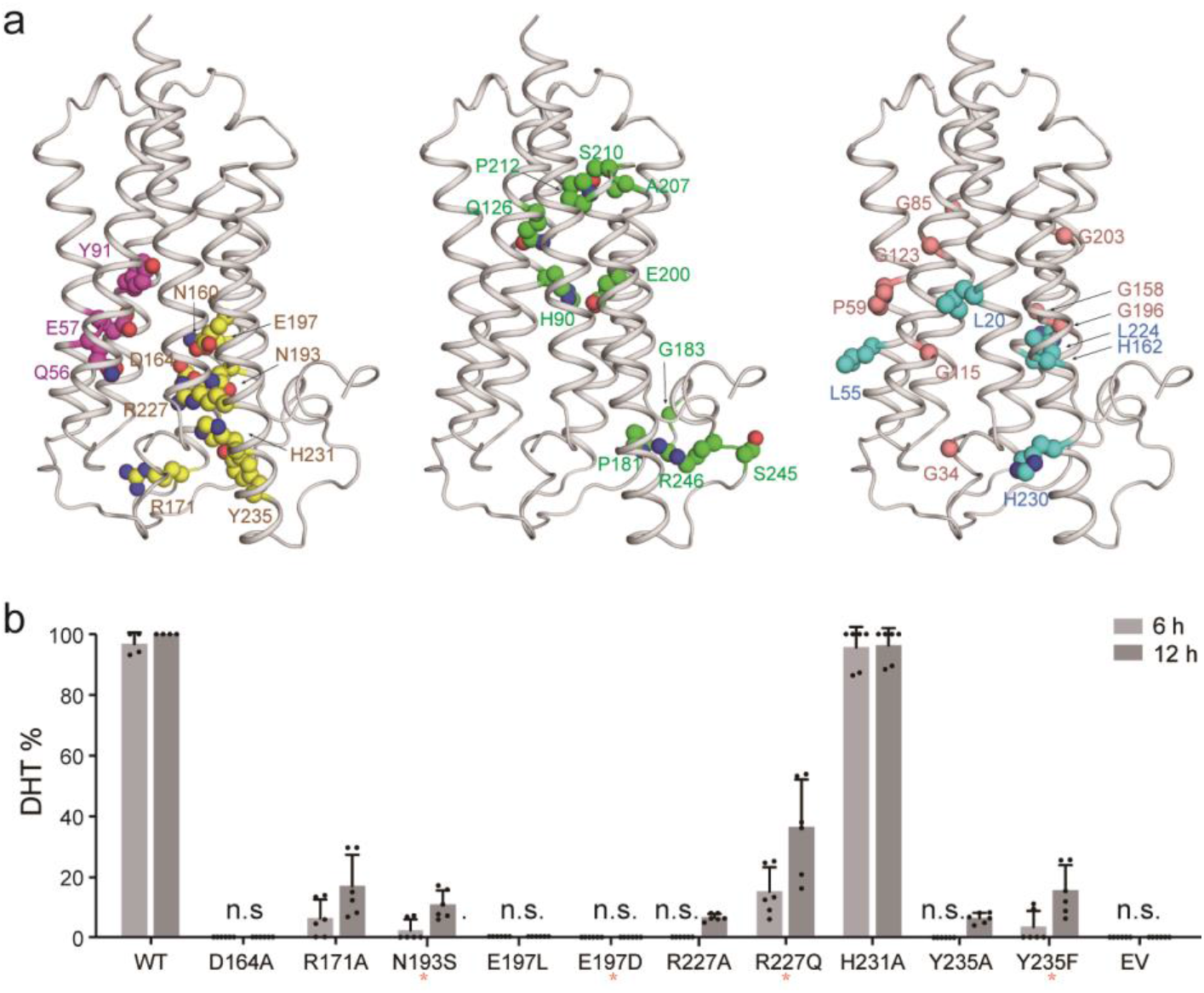
Disease related loss-of-function mutations in HsSRD5A2 model. **a,** 32 disease related loss-of-function mutations were classified in 5 categories. Catalytic site residues are in magenta, NADPH binding residues are in yellow, structural destabilizing mutations are in green, wheat or cyan based on residue properties. **b,** Biochemical analysis of disease related HsSRD5A2 variants. All variants were transient expressed in HEK293T cells for 24 hours. The cells were treated with [3H]-labeled T for 6 or 12 hours. The collected and homogenized cell membranes were treated with [^3^H]-labeled T. HsSRD5A2 activities were measured by the percentage of DHT detected by HPLC. Data are mean±s.d. derived from technically independent experiments in duplicate. Each experiment was reproduced at least three times on separate occasions with similar results. Red stars (*) below variant labels indicated the variants in the patients.

## Discussion

Enzymes of steroid 5α-reductase family, which catalyze the 3-oxo-Δ^4^ reduction reactions, have critical roles in steroidogenesis. Notably, all SRD5A enzymes share high sequence similarity from bacteria to mammal (Extended Data Fig. 8). On the basis of our structural observations and biochemical analysis, we firmly believed that PbSRD5A is a proper prototype to propose the catalytic model to shed light on the mechanistic understanding of all SRD5A enzymes.

It is not surprising that SRD5As exhibit similar fold and the NADPH cofactor-binding sites are highly conserved in the steroid reductase families. The structural and biochemical characterizations reported herein illuminated new features of steroid substrate recognition, the catalytic mechanism, and positions of natural mutations associated with steroidogenesis deficiency. Our results suggested that, if the C-3 carbonyl oxygen atoms of the substrates displace the water molecule in PbSRD5A structure, the side chain of conserved glutamate (E57) should be protonated and may serve as dual functions. On one hand, the glutamate may form a hydrogen bond to the carbonyl oxygen of substrates and, as a super acidic hydrogen bond donor, to help activate α, β-unsaturated ketone moiety of substrates for carbon-carbon bond reduction by NADPH (Extended Data Fig. 9a-c). On the other hand, except for the hydride (H^−^) provided by NADPH, one additional proton is necessary to complete the Δ^4^ carbon-carbon double bond reduction. The disease-related variant E57Q in HsSRD5A2, mimicking the constitutive protonated state, lost the reduction activity (Fig. 3f). These results indicated that the protonation of glutamate is not sufficient and this glutamate should act as the proton donor to complete the reduction reaction as well (Extended Data Fig. 9d). The structural analysis and MD simulation results also supported our speculation that, in the presence of substrate, the extra space is limited to accommodate additional water molecules in the catalytic site to provide proton.

In terms of future research, one major question awaits further investigation: the substrate and inhibitor selectivity of SRD5As. As mentioned above, PbSRD5A mainly responded to progesterone, whereas HsSRD5A1 and −2 had significant reduction activity to androstenedione and testosterone, respectively (Fig. 1b, c). Moreover, dutasteride showed higher inhibitory potency than finasteride, which is similar to SRD5A1 but not −2 (Fig. 1d) ^27^. We noticed that the cytosolic half of TM1 and −2 of all SRD5As may play critical roles in recognizing the “tail” part attached to C-17 of steroid substrates. Intriguingly, the movements of TM1s (Extended Data Fig. 7a), combined with the observation of significantly large B factor value of TM1 in PbSRD5A structure (Extended Data Fig. 1e), suggested that the TM1 may play important roles in the-substrate entering and releasing. Based on the sequence alignment over 150 SRD5As, the cytosolic half of TM1 and −4 are the most variable regions (Extended Data Fig. 6 and 8). The sequence diversity on TM1 and −4 may also contain the recognition “code” contributing to the specificity for varies of steroid substrates, as well as varies of inhibitors. Although our knowledge of these enzymes is not yet complete, the results we described here should serve as a foundation for future exploring the substrate and inhibitor selectivity, which is one of the determinants to successfully design therapeutic molecules targeting SRD5As with improved specificity and therapeutic effects.

## Methods

### Cloning and expression of PbSRD5A

The cDNA of *Proteobacteria bacterium* SRD5A (GenBank ID RIL07334.1, denoted as PbSRD5A) was synthesized from Genescript and subcloned into pFastBac vector (Invitrogen) with amino-terminal 10XHis tag. Baculovirus was generated with the Bac-to-bac system (Invitrogen) and used for infecting *Spodoptera frugiperda* (Sf9) cells (Sino-Bio) at density of 2×10^6^ cells per mL and 10 mL of virus per liter of cells. Infected cells were collected after 60 h by centrifugation, frozen in liquid nitrogen and stored at −80°C.

### Purification of PbSRD5A

Cell pellets were disrupted using Dounce tissue grinder (homogenizer) for 60 cycles on ice with lysis buffer (25 mM HEPES pH 7.5, 150 mM NaCl, 5% v/v glycerol) and subjected to ultracentrifugation at 40,000 x rpm for 60 min at 4 °C. The supernatant was discarded and the membrane fractions were homogenized in 20 mL solubilization buffer (25 mM HEPES pH 7.5, 150 mM NaCl, 5% v/v glycerol) with cocktail inhibitors (Sigma) per initial liter of culture volume. Membrane fractions was dissolved by the addition of 2% (w/v) n-dodecyl-β-D-maltopyranoside (DDM, Bluepus) and rotated at 4 °C for 2 h. Cell debris was pelleted at 18,000 x rpm for 30 min at 4 °C, and the supernatant was loaded on Ni^2+^-nitrilotriacetate affinity resin (Ni-NTA, Qiagen), 0.5 mL resin per liter cell culture. The resin was washed twice with 40 column volumes of wash buffer (25 mM HEPES pH 7.5, 150 mM NaCl, 5% v/v glycerol, 30 mM imidazole pH 8.0, 0.02% w/v DDM), and eluted with elution buffer (25 mM HEPES pH 7.5, 150 mM NaCl, 5% v/v glycerol, 250 mM imidazole pH 8.0, 0.02% w/v DDM). After concentrating, the protein was treated by 0.5 mg/mL drICE protease and incubated with 2 mM NADPH (Sigma-Aldrich) for 2 h at 4 °C. The protein was applied to size-exclusion chromatography on a Superose 6 Increase 10/300 GL column (GE healthcare) in gel filtration buffer (25 mM HEPES pH 7.5, 150 mM NaCl, 5% v/v glycerol, 0.02% w/v DDM, 5 mM DTT). Fractions containing the highest concentration of PbSRD5A were collected for crystallization.

### Lipid cubic phase crystallization

Purified PbSRD5A protein was concentrated to 25 mg/mL and incubated with another 10 mM NADPH before preparing the cubic phase. We reconstituted the protein into lipid cubic phase (LCP) by mixing with molten monoolein at a protein/lipid ratio of 2:3 (v/v) using a syringe lipid mixer. The meso-phase was dispensed in 60 nL drops onto 96-well glass plates and overlaid with 800 nL precipitant solution. The crystallization trials were performed using a Gryphon robot arm (Art Robbins Instruments). Crystals grew from buffers containing 40% PEG 400, 100 mM HEPES pH 7.5, 100 mM potassium acetate, and reach full size within one week at 20 °C.

### Initial model building of PbSRD5A

The key components of 3D model construction pipeline consist of three parts: (i) multiple MSA generation from a diverse set of sequence databases under different sequence search engines ^51^, (ii) a fused deep residual network that takes input multiple MSAs to predict inter-residue distances and orientations, and (iii) a fast Rosetta model building protocol based on restrained minimization with distance and orientation restraints ^52^. Nine alternative MSAs were generated from 3 sequence databases: UniClust30 ^53^, UniRef90 ^54^, and NCBI_NR ^55^, and for each database we used HHblits ^56^, JackHMMER, and BLAST to search for homology sequences. To fuse these 9 alternative MSAs, we implemented a deep residual network with a strip pooling module to effectively capture long-range relationship of residual pairs. Such fuse architecture could alleviate the issue caused by a problematic MSA that could decrease the prediction accuracy. Following trRosetta and AlphaFold, we generated the 3D structure model from the predicted distance and orientation using constrained minimization. Specifically, the predicted distance- and orientation-probabilities are first converted into potentials, which are then used as restraints to be fed into Rosetta together with centroid level (coarse-grained) energy optimization. Afterwards, the top 50 folded structures satisfying the restraints were selected according to Rosetta energy for further full-atom relaxation by Rosetta. Finally, 5 best models were selected from the 50 structures using GOAP energy.

### Data collection and structure determination

The X-ray diffraction data set was collected at SSRF (Shanghai Synchrotron Radiation Facility) beamlines BL18U1. The data set was further integrated and scaled with XDS Package ^57^ and Aimless in the CCP4 ^58^ suite, respectively. The structure of PbSRD5A was solved by molecular replacement (MR) with the predicted model as search model using PHASER ^59^. The structure was refined by PHENIX ^60^ and Coot ^61^. The related statistics are listed in Extended Data Table 1.

### *In vitro* enzymatic assay of PbSRD5A

Purified wild type or mutated PbSRD5A of 100 ng were incubated with [^3^H]-labeled progesterone (~500,000 cpm, 7.5 nM), progesterone (10 nM) and dutasteride (10 μM) or finasteride (10 μM) in 0.2 ml of PBS buffer with 0.02% DDM (pH 7.4) and 0.2 mM NADPH at 37 °C for 1 h. The reaction was stopped by the addition of 0.5 ml ethyl acetate: isooctane (1:1). Steroids were extracted and analyzed by HPLC. PbSRD5A *in vitro* metabolism assay for other two substrates, testosterone and androstenedione, was performed using the same protocol. All experiments have been repeated for three times and the results represent mean and standard deviation (s.d.).

### HPLC analysis

High-performance liquid chromatography (HPLC) analysis was performed on a Waters Acquity ARC HPLC (Waters, Ireland). Dried samples were reconstituted in 100 μL of 50% methanol and injected into the HPLC. Metabolites were separated on CORTECS C18 reverse-phase column 4.6 × 50 mm, 2.7 μM (Waters, Ireland), using a methanol/water gradient at 40 °C. The column effluent was analyzed using β-RAM model 3 in-line radioactivity detector (LABLOGIC, USA). Results represented the mean and standard deviation (s.d.) value from three repeated experiments. All HPLC studies were run in duplicate and repeated at least three times independently.

### Enzymatic assay of HsSRD5A2

HEK293T cells were purchased from the American Type Culture Collection (Manassas, VA) and maintained in DMEM with 10% FBS (ExCell Bio, China). All experiments were done in plates coated with poly-DL-ornithine (Sigma-Aldrich, St. Louis, MO). Cell lines were authenticated by Hybribio (Guangzhou, China) and determined to be mycoplasma free with primers 5’-GGGAGCAAACAGGATTAGATACCCT −3’ and 5’-TGCACCATCTGTCACTCTGTTAACCTC −3’. Cells were seeded and incubated in 24-well plate for 24 h before transfected with the indicated plasmids. Cells were then treated with a mixture of [^3^H]-labeled steroids (final concentration, 10 nM T and 10 nM progesterone; ~1,000,000 cpm/well; PerkinElmer, Waltham, MA) at 37 °C. Aliquots of medium were collected and treated with 300 units of β-glucuronidase (Novoprotein Scientific Inc, China) at 37 °C for 2 hours, extracted with ethylacetate:isooctane (1:1), and dried in a freeze dryer (Martin Christ Gefriertrocknungsanlagen, Germany). Steroids were extracted and analyzed by HPLC.

### Substrate docking and MD simulation for PbSRD5A, HsSRD5A1 and −2

Optimized binding poses of the pair of ligands and modeled SRD5A1/2-NADPH complexes were predicted by iteratively employing molecular docking and molecular dynamics (MD) simulations. The NADPH was initially placed in the models by structural alignment of the HsSRD5A1/2 models and PbSRD5A structure; and the testosterone was docked into the initial structure of the binary complex by AutoDock Vina ^62^. The ternary complex was embedded in a lipid bilayer constructed according to the compositions of ER membrane. The whole system was solvated in a TIP3P water with 0.15 M of NaCl, following by 200 ns of restrained MD simulations to fully relax and equilibrate the solvent and membrane structure at 303.15 K and 1 bar. Here, protonation state of the protein was assigned by the web server H++ assuming pH 7.4, and charmm36m force field was employed in simulations. Finally, 50 ns of plain MD simulation without any restraint was carried out to capture the thermodynamics effect on the conformational fluctuation of the complex. The snapshots of the protein/NADPH complex were recorded every 100 ps in our production MD, generating 500 conformers in total for each complex. Then, molecular docking was carried out for each of the 500 conformers generated from our MD simulations using AutoDock Vina. The protonation states of the substrates and the NADPH-protein complexes were kept as in the MD simulations. A 25× 25 × 25 Å3 box centered at the ring of nicotinamide group of NADPH, was considered as the searching region for each conformer. Flexibilities of the substrates and all sidechain of the residues within the searching region were considered. Exhaustiveness (80) docking was carried out to search the optimized binding pose, with the so-called Broyden-Fletcher-Goldfarb-Shanno (BFGS) algorithm for local optimization. For data analysis, all the docked conformers were sorted according to the binding affinities scored by Vina. The docked result with the strongest binding affinity among the 500 conformers was considered as the first representative binding pose. Clustering analysis was performed to group the 500 conformers, considering the coordinates of only the heavy atoms of proteins. The representative of a cluster of conformers, which had the highest binding affinity on average, was considered as the second representative binding pose. The correlation between predicted binding affinity and structural displacement of the docked ligand from the one in the first binding pose was assessed by calculating their root-mean-square distance (RMSD) for the ligand aligned according to the heavy atoms of the protein.

### Analytical ultracentrifugation analysis

Sedimentation velocity analysis was carried out with an XL-I analytical ultracentrifuge (Beckman Coulter) with An-50 Ti rotor for PbSRD5A and MvINS at 20 °C. Reaction buffer containing 25 mM HEPES pH 7.5, 150 mM NaCl, 5% v/v glycerol, 0.02% w/v DDM, and 5 mM DTT was used while the reference solution was without DDM. All data were collected at a speed of 50,000 x rpm. Samples were monitored real time using UV absorption at 280 nm and optical refraction at intervals of 4 min. Buffer along was used as reference. The molecular weights were calculated by the SEDFIT^63^ and GUSSI programs.

## Acknowledgments

We are grateful to Shanghai Synchrotron Radiation Facility (SSRF) for providing x-ray beam time and onsite assistance. We also thank Guijuan Cheng and Jie Ni at the Chinese University of Hong Kong, Shenzhen for critical discussion, figure and manuscript revision.

## Author contributions

R. R, D. D., and F. L. initiated the project. F. W. and Y. Z. aligned and ordered all genes. Y. H. and P. C. made all PbSRD5A constructs and purified the proteins. Y. H. and B. P. crystallized the protein. B. S. and Q. X. collected and analyzed all diffraction data, and determined the structure. Z. L. and S. W. provided the initial model. Y. C., W. L. and L. Z. performed docking, MD simulations based on structural and biochemical results, and energy minimization. Q. Z. performed all radio isotope HPLC analysis. R. R, Z. L. and D. D. supervised the project. R. R., D. D. and Z. L. prepared the manuscript.

## Data and materials availability

The coordinates and structure factors of the PbSRD5A protein have been deposited in the Protein Data Bank with accession code 7C83.

