## Supplementary figures for "Crystal structure of steroid reductase SRD5A reveals conserved steroid reduction mechanism"

### Extended Data

#### Extended Data Figures 1-9

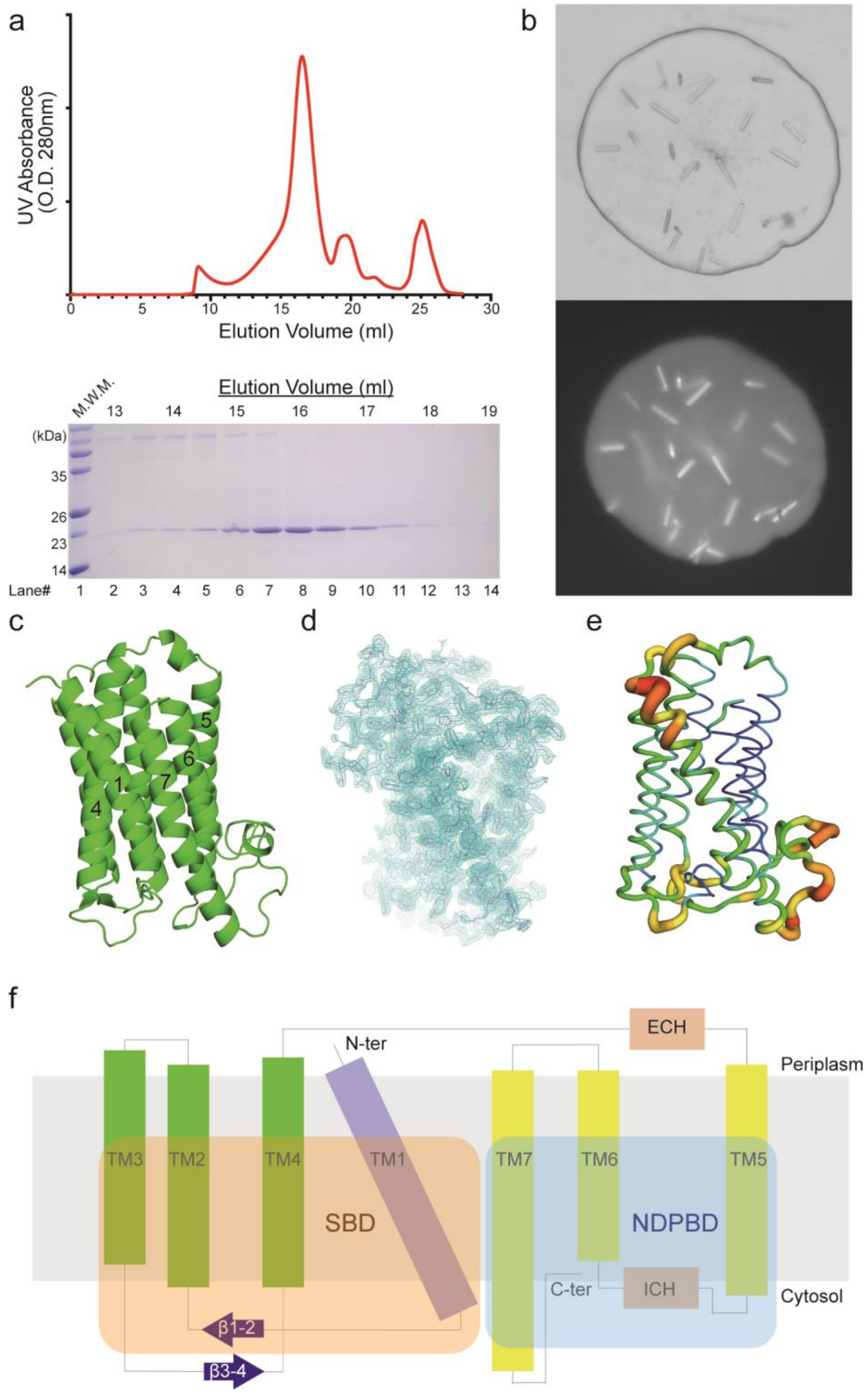


**Extended Data Fig. 1 | Purification, crystallization and structural determination of PbSRD5A. a,** The representative size exclusion chromatography of PbSRD5A and SDS-PAGE gel stained by Coomassie Brilliant Blue R-250. **b,** Photograph of PbSRD5A crystal in lipidic cubic phase under visible (upper) and UV light (lower). **c,** The initial predicted model of PbSRD5A is shown as green cartoon. **d,** The $2F_o$-$F_c$ electron density map of PbSRD5A (cyan mesh) is contoured at 1.2σ. **e,** The b factor of PbSRD5A is colored by spectrum. Red and blue represent the highest and lowest b factor values, respectively. **f,** Topology of PbSRD5A. Substrate binding domain (SBD) and NADPH binding domain (NDPBD) are shadowed in orange and cyan, respectively.

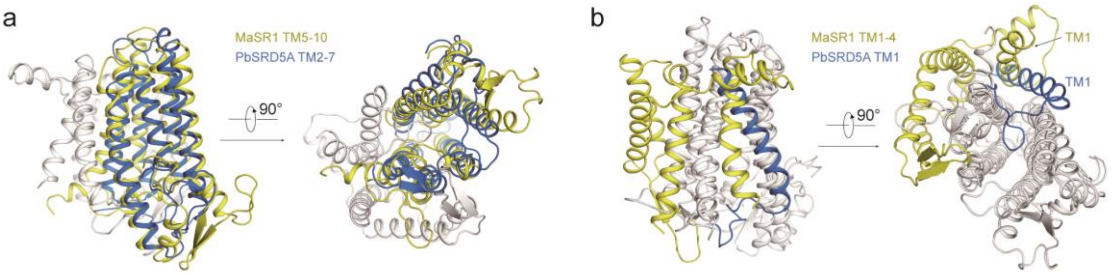


**Extended Data Fig. 2 | Structural alignment of PbSRD5A and MaSR1. a,** TM2-7 of PbSRD5A and TM5-10 are colored marine and yellow, respectively. Two perpendicular views are shown. TM2-7 of PbSRD5A and TM5-10 are superimposed with the r.m.s.d. of 3.60 Å over Cα of 184 residues. **b,** TM1 of PbSRD5A and TM1-4 are colored in marine and yellow, respectively. Two TM1s adopt distinct conformations and TM2-4 of MaSR1 is missing in PbSRD5A.

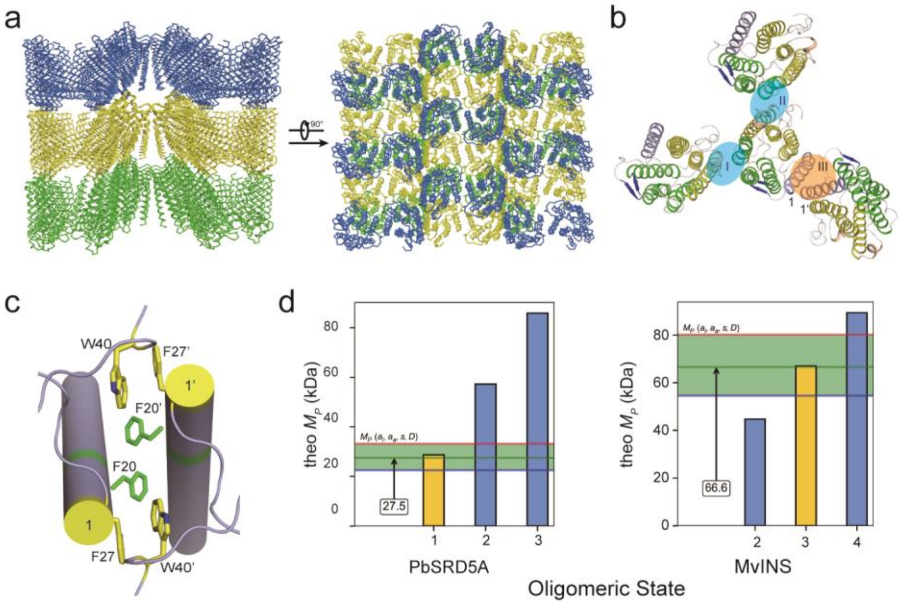


**Extended Data Fig. 3 | Crystal packing and oligomerization state examination of PbSRD5A. a,** Two perpendicular views of the crystal packing of PbSRD5A in the space group of $C222_1$. **b,** One PbSRD5A molecule interacts with three adjacent molecules in the crystal. The interface I and II are highlighted in cyan shadow and interface III is in orange. **c,** The hydrophobic residues involved in interface III are shown as yellow and green sticks. **d,** The estimated molecular weights were calculated by AUC analysis. Ranges of PbSRD5A and MvINS are shown in palegreen. X-axis values are the putative oligomerization states.

a

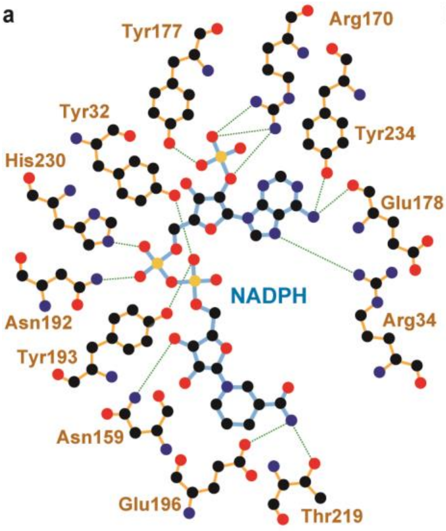


b **PbSRD5A-NADPH Hydrogen Bond Networks by PISA**

| No. | NADPH | Distance (Å) | PbSRD5A |
|---|---|---|---|
| 1 | N6A | 2.68 | Glu178 [O] |
| 2 | N6A | 3.05 | Tyr234 [OH] |
| 3 | N7A | 2.83 | Arg34 [NH2] |
| 4 | N7N | 2.98 | Glu196 [OE1] |
| 5 | N7N | 3.21 | Thr219 [OG1] |
| 6 | O1A | 2.93 | Asn192 [ND2] |
| 7 | O1N | 2.78 | Tyr32 [OH] |
| 8 | O1N | 2.48 | Tyr193[OH] |
| 9 | O1N | 3.51 | Arg226 [NE] |
| 10 | O1X | 2.60 | Tyr177 [OH] |
| 11 | O2A | 3.19 | Tyr193 [OH] |
| 12 | O2A | 2.80 | His230 [NE2] |
| 13 | O2B | 2.77 | Arg170 [NH1] |
| 14 | O2D | 3.41 | Asn159 [ND2] |
| 15 | O2X | 2.93 | Arg170 [NH2] |
| 16 | O2X | 2.76 | Arg170 [NH1] |
| 17 | O3 | 3.49 | Asn192 [ND2] |
| 18 | O3B | 3.21 | Arg99 [O] |
| 19 | O3D | 2.98 | Arg90 [NH2] |
| 20 | O3D | 2.78 | Asn159 [ND2] |
| 21 | O3D | 2.63 | Asp163 [OD2] |
| 22 | O7N | 3.51 | Asn222 [ND2] |

c **Other PbSRD5A-NADPH Interactions by LigPlot+**

| No. | PbSRD5A | Interactions | No. | PbSRD5A | NADPH | Interactions |
|---|---|---|---|---|---|---|
| 1 | Trp50 | Hydrophobic interaction | 1 | Arg34 [NE] | O1X | Water mediated hydrogen bonds |
| 2 | Phe94 | Hydrophobic interaction | 2 | Val100 [O] | O3X | Water mediated hydrogen bonds |
| 3 | Met98 | Hydrophobic interaction | 3 | Leu166 [O] | N3A | Water mediated hydrogen bonds |
| 4 | Leu166 | Hydrophobic interaction | 4 | Asn192 [O] | O1A | Water mediated hydrogen bonds |
| 5 | Leu169 | Hydrophobic interaction | 5 | Arg226 [NH2] | O2A | Water mediated hydrogen bonds |
| 6 | Leu223 | Hydrophobic interaction | | | | |

**Extended Data Fig. 4 | a,** Residues that directly form hydrogen bonds with NADPH are shown in stick-ball model. The bonds of residues and NADPH are colored in yellow and cyan, respectively. The atoms are colored by elements (black for carbon, red for oxygen, purple for nitrogen, and yellow for phosphorus). The diagram is generated by LigPlot+[1]. **b,** The hydrogen bond distances between atoms in PbSRD5A residues and NADPH are measured by PISA[2]. The residues shown in panel **a** are colored in brown. Other hydrogen bonds only found by PISA are colored in black. **c,** The indirect interactions between PbSRD5A and NADPH were listed as two categories. The hydrophobic interactions and water mediated polar interactions are listed in the left and right panels, respectively. These interactions are analyzed by LigPlot+[1].

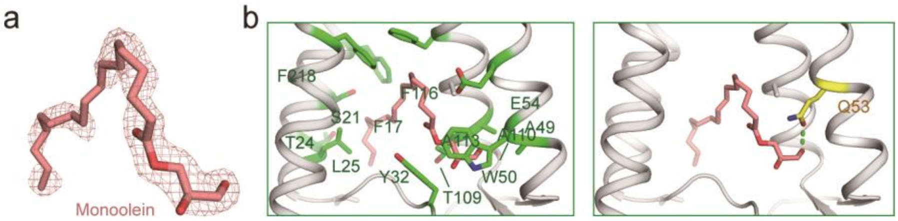


**Extended Data Fig. 5 | PbSRD5A accommodates one MAG molecule in crystal structure. a,** Monoolein fits into the density map shown in Fig. **2b**. **b,** Coordination of monoolein in PbSRD5A. The residues that interact with monoolein with hydrophobic effect are shown as green sticks. Q53, shown as yellow sticks, forms hydrogen bond (green dash) with the hydroxyl group of monoolein.

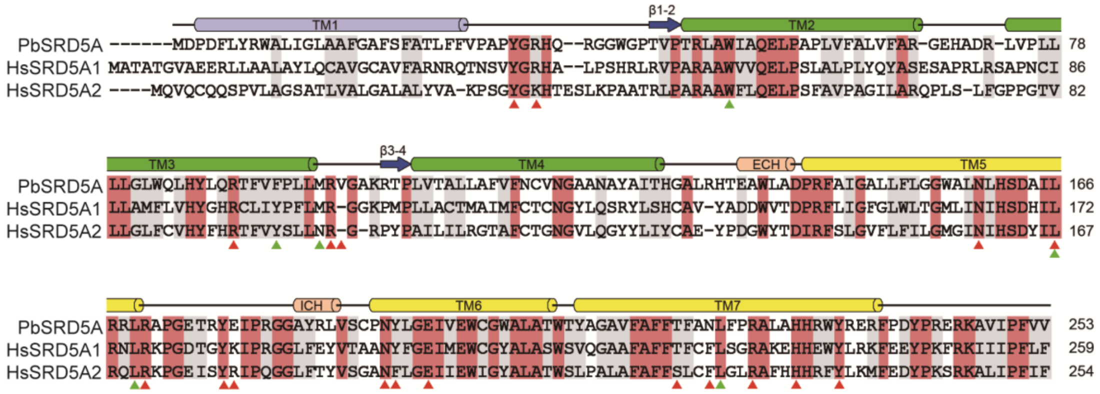


**Extended Data Fig. 6 | Sequence alignment of PbSRD5A with HsSRD5A1 and -2.** Secondary structural elements of PbSRD5A are indicated above the sequence alignment. Invariant and highly conserved amino acids are shaded in rose red and grey, respectively. The residues identified for NADPH binding are highlighted at the bottom by red (polar interactions) and green triangles (hydrophobic effects), respectively.

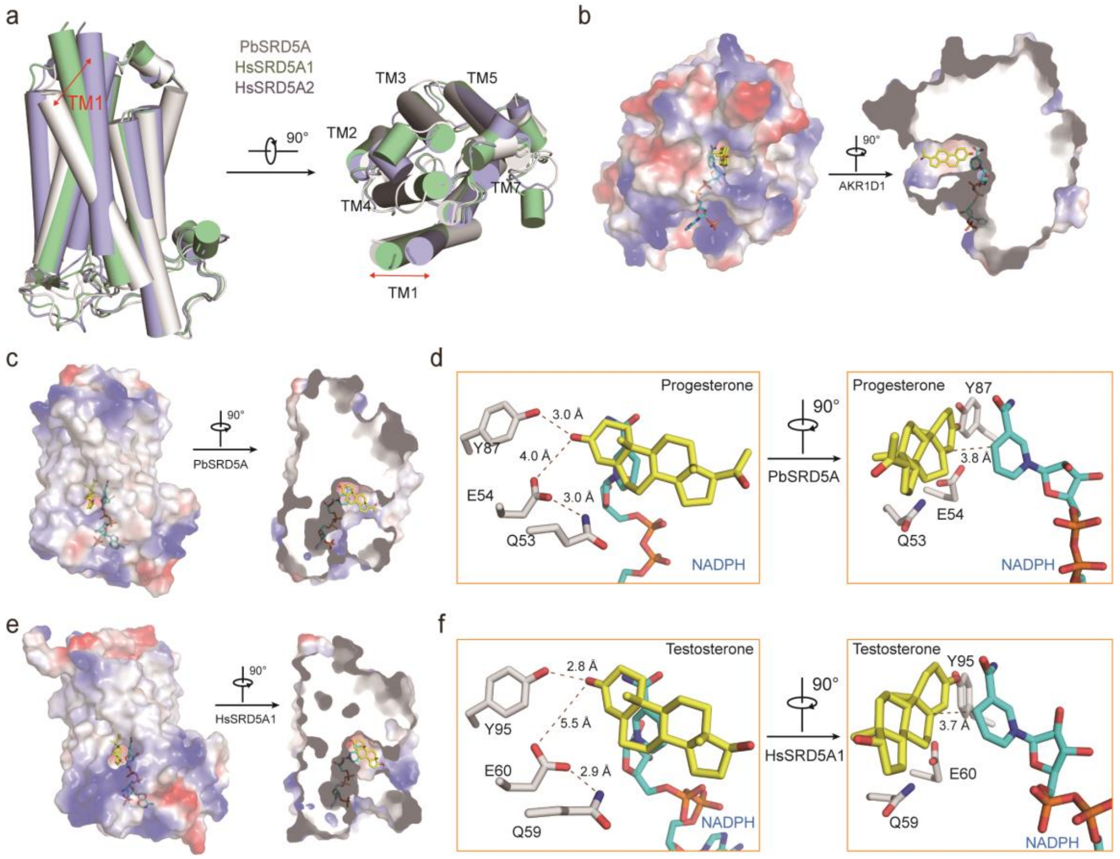


**Extended Data Fig. 7 | Structural comparison and substrate docking of SRD5As.** **a,** The superposition of PbSRD5A (gray), HsSRD5A1 (palegreen) and HsSRD5A2 (lightblue) is shown as cylindrical cartoon. The arrows indicate the conformational difference of TM1s in SRD5As by two perpendicular views. **b,** The structure of AKR1D1-progesterone (PDB code: 3COT). The semi-transparent electrostatic surface of HsSRD5A2 is shown. **c,** The docking pose of progesterone in PbSRD5A docking model. **d,** The coordination of conserved Q-E-Y motif with progesterone in PbSRD5A docking model. **e,** The docking pose of progesterone in HsSRD5A1 docking model. **f,** The coordination of conserved Q-E-Y motif in HsSRD5A1 docking model. In (**b**)-(**f**), substrates are shown as yellow stick. NADPH is colored cyan. Two perpendicular views are shown.

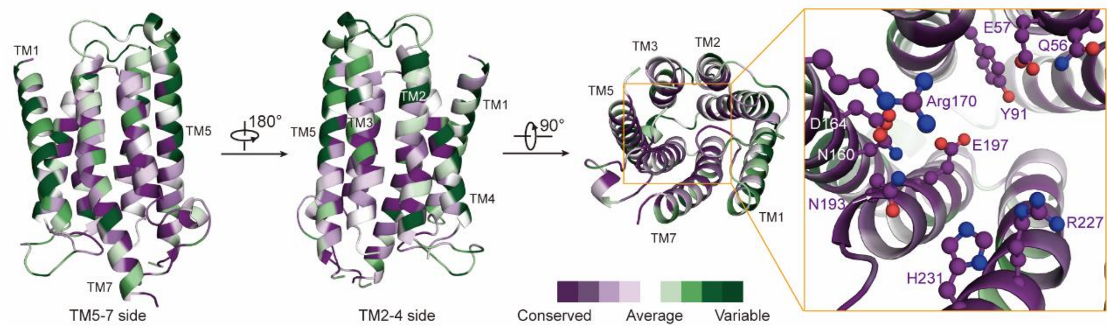


**Extended Data Fig. 8 | Mapping of the conserved residues of steroid 5-alpha reductases to the structural model of HsSRD5A2.** Sequence alignment was made for 150 steroid 5-alpha reductases and putative homologues. The conserved residues were mapped to the structural model of HsSRD5A2 using ConSurf[3,4]. Invariant residues in NADPH binding and catalytic site residues are show in stick-ball model in the inset.

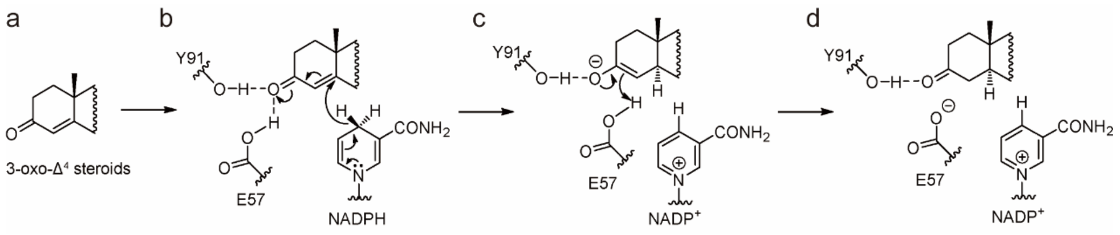


**Extended Data Fig. 9 | Reduction reaction mechanism of SRD5As in catalyzing 3-oxo-Δ$^4$ steroids. a,** Key structural features of 3-oxo-Δ$^4$ steroids. **b,** the glutamate (E57 in HsSRD5A2) may form a hydrogen bond to the carbonyl oxygen of substrates and, as a super acidic hydrogen bond donor, to help activate α, β-unsaturated ketone moiety of substrates **c,** The proposed mechanism of reduction involves a two-step reduction by the enzyme and NADPH. In the first step, NADPH coordinates on α face of the substrate and adds a hydride to C-5, leading to a selective reduction at C-5 to form an enolate ion intermediate. **d,** Secondly, the resonance-stabilized enolate ion is protonated by E57 at C-4 and thus releasing a saturated ketone product and an NADP$^+$.

### Extended Data Tables 1-2

| Protein | PbSRD5A (PDB code: XXXX) |
|---|---|
| **Integration Package** | XDS |
| Beamlines of SSRF | BL18U1 |
| Space Group | $C222_1$ |
| Unit Cell (Å) | 52.2 104.3 123.0 |
| Unit Cell (°) | 90 90 90 |
| Wavelength (Å) | 0.9793 |
| Resolution (Å) | 30-2 (2.05-2) |
| $R_{merge}$ (%) | 0.098 (0.844) |
| $R_{pim}$ (%) | 0.041 (0.347) |
| $CC_{1/2}$ [c] | 0.998 (0.804) |
| I/sigma | 12.8 (2.3) |
| Completeness (%) | 99.6 (99.3) |
| Number of measured reflections | 152510 (10988) |
| Number of unique reflections | 22993 (1628) |
| Redundancy | 6.6 (6.7) |
| $R_{work}$ / $R_{free}$ (%) | 19.34/23.21 |
| No. atoms | 2298 |
| Protein | 2008 |
| ligand | 200 |
| water | 90 |
| Average B value (Å$^2$) | 36 |
| Protein | 34.44 |
| ligand | 49.2 |
| water | 41.2 |
| R.m.s. deviations | |
| Bonds (Å) | 1.28 |
| Angle (°) | 0.02 |
| Ramachandran plot statistics (%) | |
| Most favorable | 98.80 |
| allowed | 1.2 |
| Disallowed | 0 |

**Extended Data Table 1 | Statistics of data collection and refinement for native PbSRD5A.** One crystal was used for structure determination. Values in parentheses are for the highest resolution shell. $R_{merge}=\Sigma_h\Sigma_i|I_{h,i}-I_h|/\Sigma_h\Sigma_iI_{h,i}$, where $I_h$ is the mean intensity of the i observations of symmetry related reflections of h. $R=\Sigma|F_{obs}-F_{calc}|/\Sigma F_{obs}$, where $F_{calc}$ is the calculated protein structure factor from the atomic model ($R_{free}$ was calculated with 5% of the reflections selected).

| Category | HsSRD5A2 | HsSRD5A1 | PbSRD5A |
|---|---|---|---|
| NADPH binding residue mutations | N160D | N165 | N159 |
| | D164V | D169 | D163 |
| | R171S | R176 | R170 |
| | N193S | N198 | N192 |
| | E197D | E202 | E196 |
| | R227Q | R232 | R226 |
| | H231R | H236 | H230 |
| | Y235F | Y240 | Y234 |
| Catalytic site mutations | Q56R | Q59 | Q53 |
| | E57Q | E60 | E54 |
| | Y91D | Y95 | Y87 |
| Structural destabilizing mutations | A207D | A212 | A206 |
| | S210F | S215 | T209 |
| | P212R | Q217 | A211 |
| | Q126R | Q131 | N124 |
| | E200K | E205 | E199 |
| | P181L | P186 | P180 |
| | G183S | G188 | G182 |
| | S245Y | F250 | E244 |
| | R246Q/W | R251 | R245 |
| Helix breaking mutations | L20P | V23 | S21 |
| | L55Q/P | V58 | A52 |
| | H162P | H167 | H161 |
| | L224P | L229 | L223 |
| | H230P | E235 | H230 |
| Small to bulky residue (Destabilizing mutations) | G34R/W | G40 | G33 |
| | P59R | P62 | P56 |
| | G85D | A89 | G81 |
| | G115D | A120 | A113 |
| | G123R | G128 | G121 |
| | G158R | L163 | A157 |
| | G196S | G201 | G195 |
| | G203S | G208 | G202 |

**Extended Data Table 2 | Disease related loss-of-function mutations in HsSRD5A2.** Disease related loss-of-function mutations in HsSRD5A2 and the corresponding residues in HsSRD5A1 and PbSRD5A are listed. The invariant residues and highly conserved residues between HsSRD5A2 and PbSRD5A are colored in red and purple, respectively.
